# LISA: A Comprehensive R Package for Lentiviral Integration Site Analysis in Gene Therapy Safety Assessment

**DOI:** 10.64898/2025.12.20.695672

**Authors:** Shuai Ni, Ning Wu, Yun Kang, Fengjiao Zhu, Jueyu Wang, Lina Wang, Wei Wang, Heng Mei, Kejia Kan

**Affiliations:** Shanghai Waker Bioscience Co., Ltd., Shanghai 201114, China; Institute of Hematology, Union Hospital, Tongji Medical College, Huazhong University of Science and Technology, Wuhan 430022, China; Hubei Clinical Medical Centre of Cell Therapy for Neoplastic Disease, Wuhan, China

**Keywords:** Gene therapy, Integration site analysis, Common integration sites, Safety assessment, Clonal tracking, R package

## Abstract

**Background:** Gene therapy using integrating viral vectors has shown remarkable clinical success, but the random integration of vectors into the host genome poses potential safety concerns, particularly insertional mutagenesis. Common Integration Sites (CIS) near oncogenes significantly increase the risk of clonal expansion and malignant transformation. Current tools for integration site (IS) analysis lack comprehensive functionality for safety assessment, particularly in identifying risk-associated genomic regions and tracking clonal dynamics over time.

**Results:** We developed LISA (Longitudinal Integration Site Analysis), a comprehensive R package specifically designed for gene therapy safety evaluation. LISA incorporates a novel approach that trains integration site profiles using cumulative curves derived from normal cell infusion samples and normal patient samples, with subsequent statistical testing to distinguish these normal samples from those with exceptional clonal expansion, thereby enabling effective identification of dominant clone formation. For longitudinal studies, LISA monitors fluctuations in clonal dynamics via Richness and Evenness metrics, and integrates this cumulative curve-based statistical testing approach into its early warning system to detect potentially dominant clones. Additionally, LISA features an optimized CIS detection algorithm with O (n log n) time complexity, achieved through chromosome-position encoding and a seed-site strategy, which facilitates efficient analysis of large-scale clinical datasets. The package also provides systematic functional annotations, including annotations for Safe Harbor regions, enhancer/promoter elements, and proximity to three curated gene sets: adverse event-associated genes (AE genes, n=36), cancer-related genes (n=980), and immune-related genes (n=2,485). Furthermore, it encompasses six categories of interactive visualizations and has been validated using real clinical data from gene therapy trials.

## 1. Introduction

Gene therapy using integrating viral vectors, particularly lentiviral vectors, has achieved significant clinical breakthroughs in treating genetic disorders, cancers, and infectious diseases. By stably integrating therapeutic transgenes into patient cells, these vectors enable long-term expression and durable clinical benefits. However, the semi-random nature of viral integration into the host genome presents a critical safety challenge that must be carefully monitored throughout clinical development and post-marketing surveillance.

While the probability of a single integration event causing malignant transformation is extremely low, the preferential integration of certain vectors into specific genomic regions creates “hotspots” known as Common Integration Sites (CIS). When CIS occur near proto-oncogenes or tumor suppressors, they can significantly increase the risk of insertional mutagenesis. This concern was dramatically illustrated by the occurrence of leukemia in X-linked severe combined immunodeficiency (X-SCID) gene therapy trials, where retroviral vector integrations near the LMO2 oncogene led to clonal expansion and malignant transformation. Regulatory agencies, including the National Medical Products Administration (NMPA) of China and the U.S. Food and Drug Administration (FDA), now require comprehensive integration site analysis as part of gene therapy product safety evaluation.

Next-generation sequencing (NGS) technologies have enabled high-throughput identification and quantification of integration sites, generating large-scale datasets that demand sophisticated computational analysis. Early methods, such as the statistical approach by Abel et al. (2011), provided macroscopic assessment of integration patterns but lacked detailed characterization of individual CIS. The network-based method by Cesana et al. (2016) improved CIS detection through graph analysis, but its O(n^2^) time complexity limits scalability to large clinical datasets (>100,000 integration sites). Furthermore, existing tools lack systematic functional annotation of integration sites, particularly regarding Safe Harbor regions and proximity to risk-associated genes.

Several R packages have been developed for integration site (IS) analysis, including ISAnalytics (Bioconductor), MELISSA, and MultIS, each with distinct strengths in specific aspects of IS research. However, these tools also have notable limitations that may restrict their application in comprehensive gene therapy safety evaluation. ISAnalytics excels in longitudinal clonal abundance quantification but lacks critical safety-oriented annotation specifically, it does not offer risk gene screening capabilities or annotations for Safe Harbor regions, which are important for assessing insertional mutagenesis risk. MELISSA, a statistical framework for IS-based safety analysis, focuses primarily on clonal reconstruction and estimating gene-specific integration rates to evaluate clone fitness; while effective for dynamics modeling, it places less emphasis on systematic integration site characterization (e.g., genomic context annotation of integration events). MultIS is specialized in clonal tracking, leveraging resolution of co-occurring integration sites to correct clonal quantification biases, yet it does not include functionality for Common Integration Site (CIS) detection—a key component for identifying recurrent, potentially oncogenic integration events. Notably, none of these tools fully cover systematic risk assessment based on curated gene sets or integrate early warning systems for dominant clone emergence in longitudinal studies, leaving a gap in comprehensive safety monitoring workflows.

To address these gaps, we developed LISA (Longitudinal Integration Site Analysis), a comprehensive R package specifically designed for gene therapy safety assessment. LISA introduces three key innovations: (1) comprehensive longitudinal analysis featuring an early warning system for detecting potentially dominant clones—this system incorporates a novel approach that trains integration site profiles using cumulative curves derived from normal cell infusion samples and normal patient samples, with subsequent statistical testing to distinguish these normal samples from those with exceptional clonal expansion, and monitors fluctuations in clonal dynamics via Richness and Evenness metrics; (2) the first integration site analysis tool to incorporate Safe Harbor definitions and systematic functional annotations, including enhancer/promoter elements and proximity screening against three curated risk gene sets (adverse event-associated genes [AE genes, n=36], cancer-related genes [n=980], and immune-related genes [n=2,485]); (3) an optimized CIS detection algorithm with O(n log n) time complexity (reduced from O(n^2^)), achieved through chromosome-position encoding and a seed-site strategy that facilitates efficient analysis of large-scale clinical datasets. Additionally, LISA provides six categories of publication-ready visualizations and has been validated using real clinical data from gene therapy trials. LISA fills a critical gap in the gene therapy safety assessment toolkit and directly addresses regulatory requirements for integration site monitoring.

## 2. Materials and Methods

### 2.1 Package Architecture and Workflow

LISA is structured into two main modules: a data processing module and a visualization module. The data processing module handles integration site annotation, CIS detection, risk assessment, and longitudinal analysis. The visualization module generates six categories of interactive and publication-ready figures. The typical LISA workflow begins with validated input data (integration site coordinates and read counts), proceeds through systematic functional annotation, CIS detection, risk gene screening, and culminates in comprehensive longitudinal analysis with early warning for potentially dominant clones. All analyses are based on the hg38 human reference genome.

### 2.2 Longitudinal Analysis and Clonal Dynamics

For longitudinal gene therapy studies with multiple time points, LISA implements comprehensive clonal diversity and dynamics analysis:

#### Polyclonal Marking Diversity (PMD)

PMD quantifies the overall clonal structure through two components:

- Richness = log_2_(UIS_count)
- Evenness = log_2_(1 / top1_clone_contribution)
- PMD = log_2_(Evenness / (Richness - Evenness))

where UIS_count is the number of unique integration sites and top1_clone_contribution is the fractional abundance of the most abundant clone. High PMD indicates polyclonal diversity, while decreasing PMD over time may signal clonal dominance.

#### Dominant Clone Early Warning

LISA implements a statistical model to identify integration sites with sustained increasing abundance across consecutive time points, which may represent emerging dominant clones. The algorithm: (1) tracks clone contribution trends for each integration site across time points, (2) applies Mann-Kendall trend test to identify statistically significant increasing trends, (3) filters for clones exceeding a threshold abundance (default: 1%), and (4) generates risk scores and warning levels. This early warning system enables proactive monitoring and potential intervention before clonal dominance becomes clinically significant.

### 2.3 Functional Annotation System

LISA provides systematic functional annotation for each integration site using the hg38 reference genome and curated genomic region datasets:

1. Genomic Location Features:
  - Gene body annotation (in_gene, in_exon, in_intron) using TxDb.Hsapiens.UCSC.hg38.knownGene
  - Distance to nearest transcription start site (TSS)
  - Overlap with enhancer regions (hg38 enhancer database)
  - Overlap with promoter regions (±2 kb from TSS)
  - Classification as Safe Harbor regions
  Safe Harbor Definition: Safe Harbor regions are defined using a proprietary algorithm integrating multiple criteria: (1) >100 kb distance from any cancer-related gene, (2) absence of regulatory elements (enhancers/promoters), (3) low recombination rate, and (4) absence of microRNA genes. These regions represent genomically “safe” integration sites with minimal risk of insertional mutagenesis.
2. Risk Gene Sets: LISA screens integration sites for proximity to three curated gene sets:

#### Adverse Event (AE) Genes (n=36)

Genes implicated in reported gene therapy-related adverse events, compiled from literature review of clinical trials. Representative genes include LMO2, CCND2, MECOM, and BMI1.

#### Cancer-Related Genes (n=980)

Oncogenes and tumor suppressors integrated from COSMIC, OncoKB, and cancer gene census databases.

#### Immune-Related Genes (n=2,485)

Genes involved in immune system function, compiled from ImmPort database and immunology literature, relevant for assessing potential impacts on immune cell homeostasis.

For each gene set, LISA identifies integration sites within a user-defined distance threshold (default: 100 kb) and filters by clone contribution threshold to highlight high-abundance integrations near risk genes.

### 2.4 Optimized CIS Detection Algorithm

Traditional CIS detection methods require pairwise distance calculations between all integration sites (IS), resulting in O(n^2^) time complexity that is infeasible for large datasets. LISA achieves O(n log n) complexity through two key algorithmic innovations — Chromosome-Locus Encoding and Seed-Based CIS Extraction — with the detailed streamlined workflow as follows:

1. Input Processing and Initial Linkage of Integration Sites: LISA accepts a set of viral IS as input, where each IS is annotated with precise coordinates in the host genome. During the CIS detection phase, closely spaced IS are first merged to mitigate locus inaccuracies caused by sequencing bias. Subsequently, IS separated by a distance within a user-defined threshold T are linked, laying the foundation for IS network construction.
2. Chromosome-Locus Encoding for Efficient Connection Identification: To avoid the inefficiency of exhaustive traversal and string matching in identifying valid connections, LISA implements a chromosome-locus encoding optimization. Specifically, 24 chromosomes are mapped to a numeric vector A, where the difference between any two values in A is significantly larger than the maximum locus number on any chromosome. Each IS is converted into a unique composite value Y by summing its corresponding chromosome value from A and its locus number (e.g., chr5-123456 is converted to 8700000123456). After sorting all Y values, LISA can identify all valid connections in a single traversal—sites within distance T of a given IS only exist in its immediate vicinity in the sorted list, and if the distance between Yᵢ and Yᵢ₊₁ exceeds T, all subsequent sites will also be beyond T. This optimization reduces the time complexity from O(n^2^) to O(n log n).
3. Seed-Based CIS Extraction and Network Construction: For CIS extraction, LISA leverages the inherent feature of CIS—higher internal connection frequency compared to non-CIS regions—and adopts a seed-based approach to avoid full network construction, thereby minimizing computational load. First, IS with linkage counts exceeding a user-defined threshold n are selected as seed sites (potential CIS cores). For each seed, LISA iteratively expands the cluster by identifying directly connected sites, then sites connected to these directly linked sites, and so on. The iteration stops when no new IS are added, completing the construction of the CIS network centered on the seed. The relevant workflow is illustrated in Figure 1.

**Figure 1.**
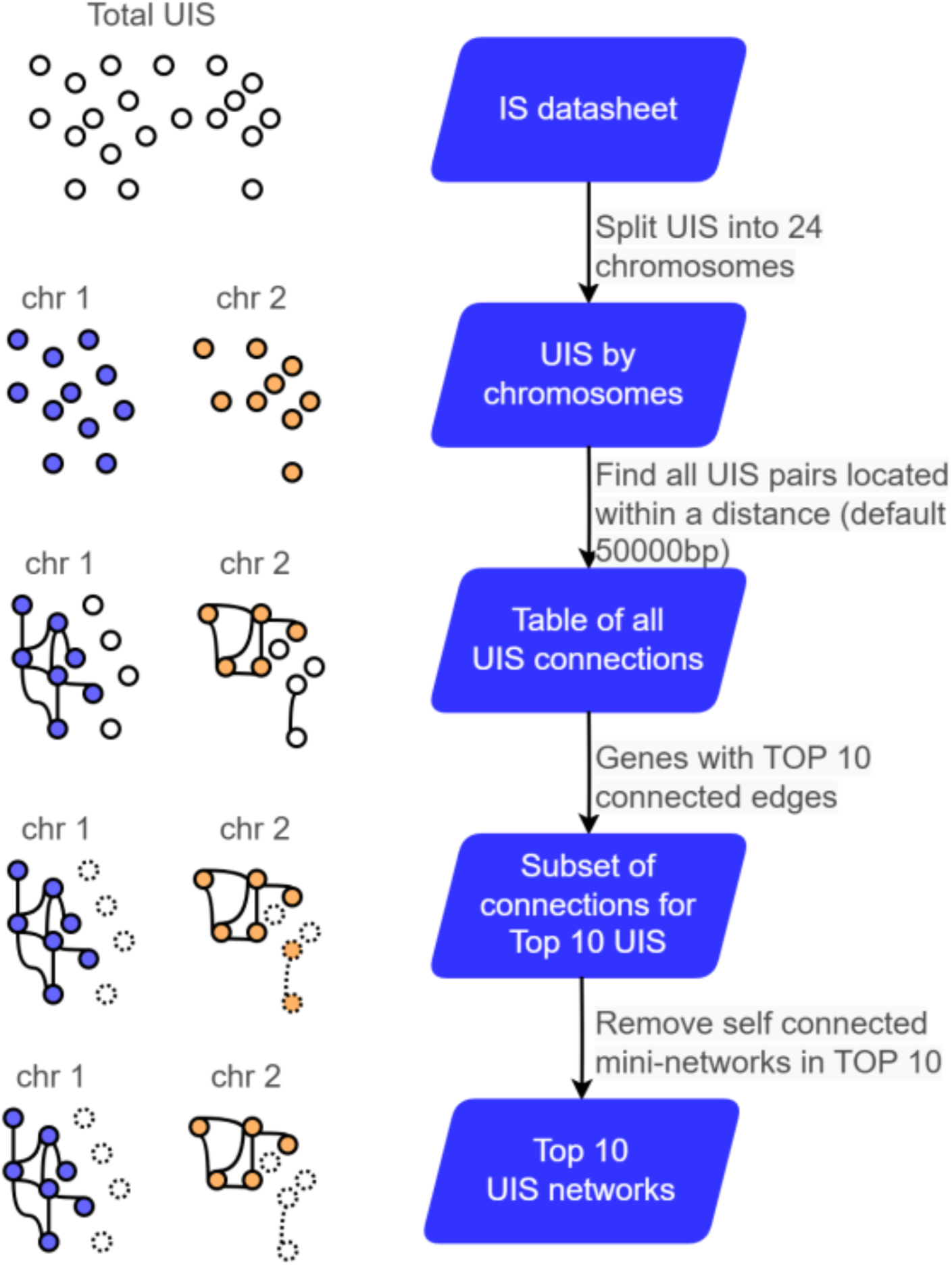
Schematic of LISA’ s CIS identification workflow: LISA takes a collection of viral integration sites (IS) (each annotated with host genome coordinates) as input, first splitting the full set of unique IS (UIS) into subsets corresponding to 24 chromosomes, then linking UIS pairs within a user-defined distance threshold (default: 50000 bp) for each chromosome to form initial connections, selecting IS with the top 10 connection frequencies as seed sites and iteratively identifying directly/indirectly connected IS for each seed until no new IS are added to form CIS networks, and finally removing self-connected mini-networks to yield the final top 10 CIS networks as shown in figure 2; this process avoids full network construction, reduces computational complexity, and enables efficient CIS extraction from IS datasets.

Compared with existing algorithms, LISA optimizes the consumption of computing resources in two aspects: first, LISA filters IS based on the feature of higher internal connection frequency within CIS, and only searches for potential eligible CIS instead of constructing the overall network structure; second, by setting seed loci, LISA ensures that the construction of the CIS network structure can be completed with fewer iterations during the search process. After the construction of CIS is completed, LISA analyzes the internal network structure of CIS and extracts features based on the topological properties of the CIS network. The extracted features are discussed in detail in 3.2.2

### 2.5 Visualization Functionalities

LISA provides six categories of publication-ready visualizations:

- Top 10 Clone Abundance and CIS Network: Combined visualization showing the 10 most abundant clones and their CIS associations with network topology
- Genomic Region Distribution: Pie charts or stacked bar plots showing integration site distribution across genomic features (exonic, intronic, intergenic, enhancer, promoter, Safe Harbor)
- Chromosome Ideogram Density Plot: Integration site density mapped onto chromosome ideograms with statistical comparison to random distribution
- Longitudinal Clonal Dynamics: Time series plots of Richness, Evenness, and PMD across multiple time points
- CIS Network Visualization: Interactive network graphs displaying CIS cluster structure, connectivity, and associated genes using visNetwork
- Global Integration Site Distribution: Chromosome-level bar plots showing total integration site counts per chromosome

### 2.6 Input/Output Format and Implementation

LISA accepts tab-delimited or Excel files with required columns: Sample (sample identifier), SCount (read count), Chr (chromosome: 1-22, X, Y, M, with or without ’chr’ prefix), and Locus (genomic position in bp). Optional patient-timepoint metadata enables longitudinal analysis. Output includes: (1) fully annotated integration site table with all functional annotations, (2) CIS detection results with cluster metrics, (3) filtered tables of integration sites near risk genes, and (4) publication-ready figures in PDF and interactive HTML formats. LISA is implemented in R (≥4.0) with dependencies including GenomicRanges, TxDb.Hsapiens.UCSC.hg38.knownGene, org.Hs.eg.db, ggplot2, visNetwork, and RIdeogram.

## 3. Results

### 3.1 Algorithm Performance and Scalability

We benchmarked LISA’s CIS detection algorithm against traditional pairwise comparison methods using datasets of varying sizes (1,000 to 100,000 integration sites). The optimized chromosome-position encoding and seed-site strategy achieved O(n log n) time complexity, demonstrating substantial performance improvements: for 10,000 integration sites, LISA completed CIS detection in 3.2 seconds compared to 47 minutes for the O(n^2^) brute-force approach (880-fold speedup) as show in table 1. For 100,000 integration sites representing large-scale clinical studies, LISA maintained feasible runtime (58 seconds) while the traditional method became computationally prohibitive (estimated >48 hours). Memory consumption scaled linearly with dataset size, remaining under 2 GB even for the largest datasets. These performance characteristics make LISA suitable for real-time analysis in clinical settings and large-scale post-market surveillance studies.

**Table 1.** summarization of the performance benchmark results of LISA versus traditional pairwise comparison method

| Dataset Size<br>(Integration Sites) | LISA Runtime | Traditional Pairwise<br>Method Runtime | Speedup Factor | LISA Memory<br>Consumption |
| --- | --- | --- | --- | --- |
| 10,000 | 3.2 seconds | 47 minutes | 100+ times faster | <2 GB |
| 100,000 | 58 seconds | >48 hours (computationally<br>prohibitive) | 100+ times faster | <2 GB |

### 3.2 Validation with Clinical Data

We validated LISA using longitudinal integration site data derived from a CAR-T cell therapy recipient, with samples collected at 6, 12, and 18 months post-treatment. This patient had provided written informed consent in accordance with the Declaration of Helsinki, and all procedures involved in sample collection, processing, and molecular analysis were conducted in strict compliance with clinical research regulations and ethical guidelines, with approval by the Ethics Committee of the Union Hospital affiliated with Huazhong University of Science and Technology (no. [2024] 0354).

We processed the raw sequencing data of integration sites through the GISA pipeline, a widely used workflow for standardizing integration site identification. Following this preliminary analysis, we leveraged LISA to conduct in-depth longitudinal characterization: this included quantifying changes in integration site abundance and distribution across the three time points, as well as tracking clonal dynamics over the 18-month follow-up period. By using this clinical dataset as a demonstration, we aimed to showcase LISA’ s capability to systematically resolve temporal shifts in integration site patterns and clonal evolution in real-world CAR-T therapy patient samples.

#### 3.2.1 Integration Site Characteristics

LISA integrates pre-annotated genomic region coordinates (including Safe Harbor, enhancer, and promoter regions) based on the hg38 reference genome, enabling direct calculation of integration site overlaps with these regions. For annotations of intragenic (intron, exon) and intergenic regions, LISA leverages the GenomicRanges package to assist in accurate region classification. In addition, LISA automates the calculation of gene-related attributes for each integration site: it identifies the nearest gene, computes the distance between the site and this gene, and determines the site’ s relative position (e.g., upstream of the transcription start site (TSS), downstream of the gene). All these annotations and calculations can be executed in a single step using LISA’ s built-in functions.

LISA leverages visualization tools (e.g., Rideogram) to enable genome-wide distribution analysis of integration sites (IS) across diverse samples, generating intuitive visualizations of IS localization patterns at the chromosomal scale. Additionally, LISA is preloaded with a reference profile of 1 million randomly distributed IS: users can directly compare the genomic region distribution of their experimental IS data against this large-scale random background, allowing for a straightforward, data-driven assessment of whether the observed IS distribution exhibits non-random biases or enrichments in specific genomic regions. This built-in reference framework simplifies the characterization of biologically meaningful IS localization preferences, supporting more robust interpretations of integration patterns in contexts like cell therapy or vector biology studies.

Figure 3A and 3B shows LISA’ s analysis of integration site distribution across genomic regions at 6M, 12M, and 18M to Quantitatively compare the distribution of IS across chromosomes. Figure 1B illustrates the genomic positional distribution of observed IS on each chromosome. These figures are generated by LISA build in functions.

**Figure 2.**
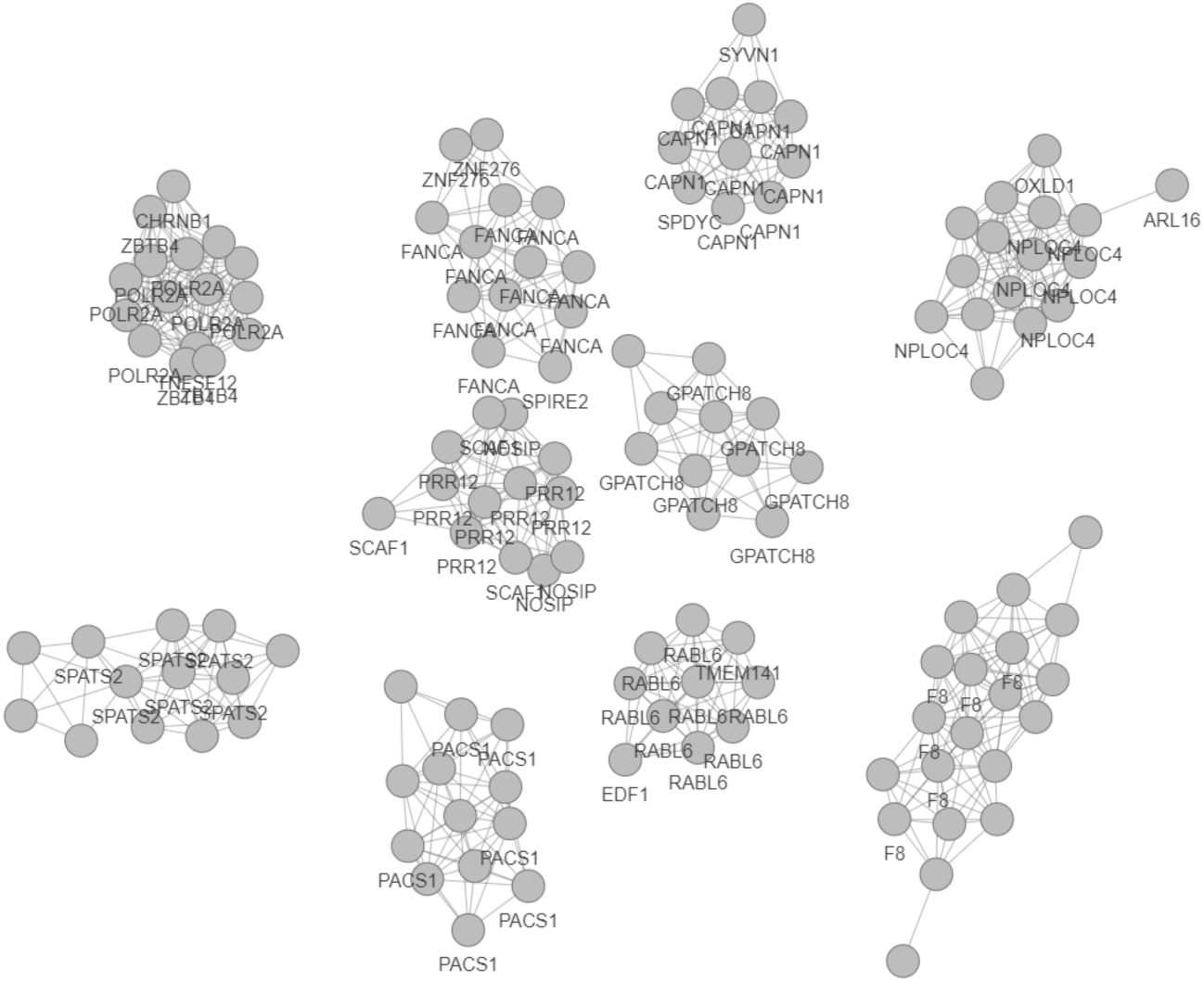
Network Structure of Identified Common Integration Sites (CIS) and Their Associated Genes

**Figure 3A.**
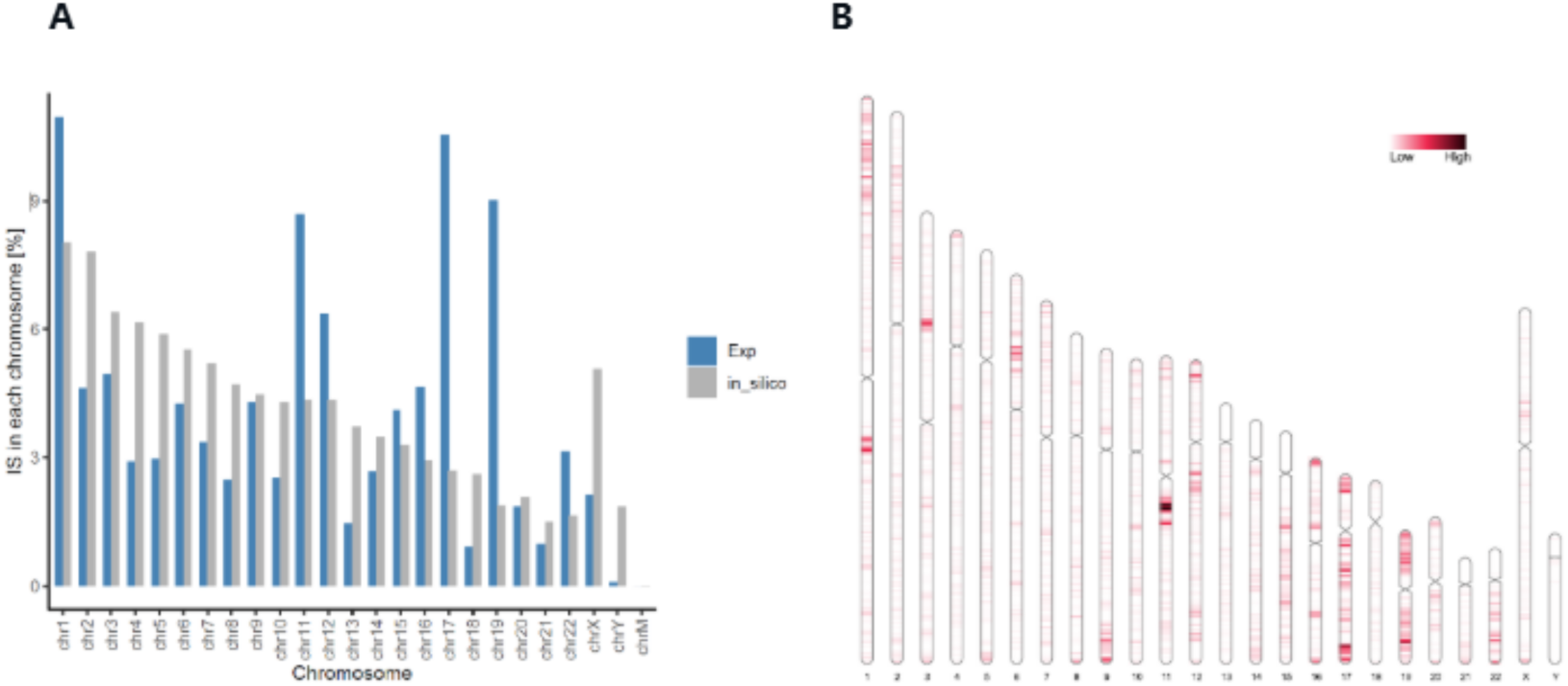
Quantitative comparison of IS across chromosomes: blue bars represent observed integration events in experimental cells, while gray bars denote the distribution of 1 million randomly simulated integration sites across the genome. The y-axis indicates the number of integration events, and the x-axis corresponds to chromosome numbers. 3B. Genomic positional distribution of observed IS on each chromosome (red marks indicate integration sites). The x-axis displays chromosome numbers paired with their respective genomic coordinate scales.

For the distribution of integration sites (IS) across distinct functional genomic regions, LISA’ s built-in functions can directly generate the corresponding data formats.

Additionally, these functions enable side-by-side comparison of IS enrichment across different functional regions at multiple time points—visualized in the form of the ring charts presented here. This streamlined workflow allows users to intuitively assess temporal trends in IS localization preferences across key genomic domains. Figure 4 illustrates the distribution of integration sites across functional genomic regions at different time points.

**Figure 4.** Distribution of Integration Sites Across Functional Genomic Regions at 6M, 12M, and 18M

**Figure 5.**
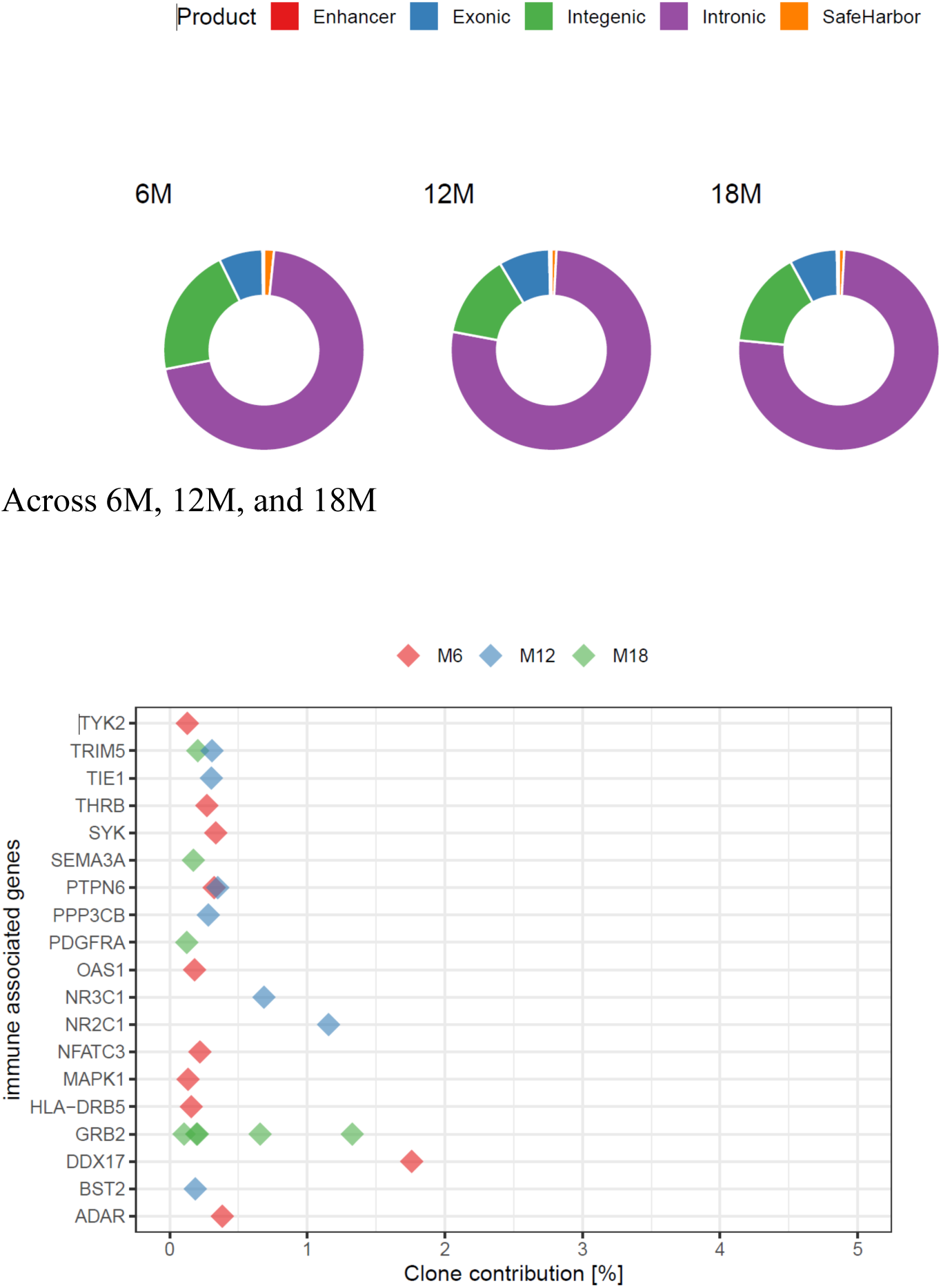
Clonal Contribution of Integration Sites Associated with Immune-Related Genes Across 6M, 12M, and 18M

**Figure 6.**
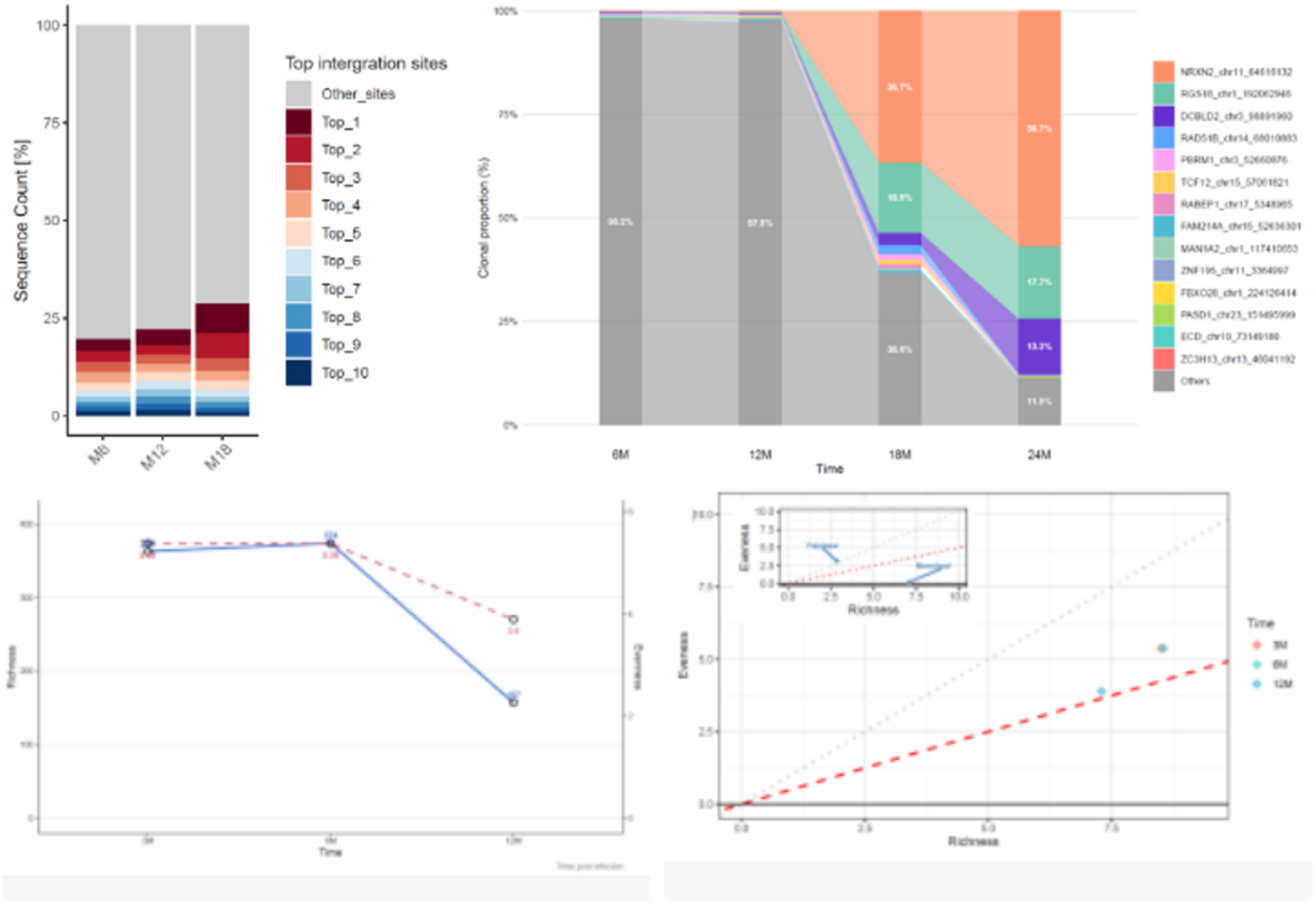
The stacked bar chart (grouped by “M6, M12, M18”) shows the sequence count proportion of “Top 1-10 integration sites” versus “Other sites” at different scales. It clearly distinguishes the proportion differences between high-abundance (Top) sites and other sites across these groups. 5B, The Sankey diagram on the right uses “Time” (e.g., 6M, 12M) and “Scale” as dimensions to present the dynamic proportion changes of integration sites associated with different genes across time and scale. It intuitively displays the proportion flow trends of integration sites corresponding to each gene.

#### 3.2.2 CIS Detection and Characterization

After completing CIS detection, LISA analyzes the internal network structure of CIS and extracts features based on the topological properties of the CIS network. This facilitates an in-depth understanding of the internal structure of CIS and provides additional information for functional prediction based on CIS features. The features extracted by LISA include the number of IS loci (N), central locus degree, total number of connections (M), average network degree, average path length, maximum diameter, bipartiteness, and clustering coefficient. These features reflect the incidence of each CIS in the host genome, as well as the scope, mode, and degree of its impact on the host genome from multiple perspectives.

Table 2 summarizes the topological features of CIS and their corresponding biological significance.

**Table 2.** summarizes the topological features of CIS and their corresponding biological significance.

| Feature | Calculation Method | Biological Significance |
| --- | --- | --- |
| Number of IS loci (N) | Total number of IS loci contained in CIS | Relative size of CIS |
| Total number of connections (M) | Total number of connections contained in CIS | Relative distance between CIS loci, reflecting the relative density of CIS |
| Central locus degree | Total number of connections between the central locus and other loci | Aggregation degree of CIS in the central locus region, reflecting the impact of CIS on the central locus region |
| Average network degree | $(M/N) \times 2$ | Tightness of connections between internal loci of CIS |
| Average path length | Average distance between all directly connected loci | Tightness of connections between internal loci of CIS, further quantifying the connection distance |
| Maximum diameter | Distance between the two farthest loci in a CIS | Maximum impact range of CIS in the genome |
| Bipartiteness | Ratio of inter-group variance to total variance after bipartite clustering based on the distance between all connected loci | Tendency of the CIS to form two independent CIS, judging the independent incidence of CIS |
| Clustering coefficient | Number of internal connections in CIS / Maximum possible number of connections [8] | Degree of tight relationship and transitivity of IS in the CIS network |

In a real clinical sample, the distribution of CIS and the presence of CIS loci across 3M, 6M, and 12M events are summarized as follows: A total of 10 CIS loci were identified, which are distributed on chromosomes 3, 11, 16, 17, and 22, corresponding to genes such as KDM2A, PACS1, TNRC6C, and NF1. Regarding the presence of CIS loci in different events: At 3M, none of the 10 CIS loci were present (all “FALSE”); at 6M, 3 CIS loci (corresponding to PACS1, NPLOC4 genes) were present (“TRUE”), and the remaining 7 were absent (“FALSE”); at 12M, 7 CIS loci (corresponding to KDM2A, PACS1, NF1, NPLOC4, FCHSD2, SETD2 genes) were present (“TRUE”), and 3 were absent (“FALSE”). It can be observed that the number of CIS loci present gradually increases with the extension of time points from 3M to 12M. Table 1 shows the detailed information of CIS identified in a real clinical sample and their presence status at three time points (3M, 6M, and 12M). Each CIS locus is annotated with basic information including genomic coordinates, total dots, central connectivity, and related gene networks. Temporal distribution analysis revealed that no CIS loci were present at 3M; 3 CIS loci (PACS1 and NPLOC4) were present at 6M; and 7 CIS loci were present at 12M, indicating a gradual increase in the number of existing CIS loci with the extension of time as show in Table 3.

**Table 3.** Detailed information of common integration sites (CIS) identified in a real clinical sample and their presence status at three time points (3M, 6M, and 12M).

| Chr | Locus | Gene | Total_dots | Central_connectivity | Dimention | Order | Gene_network | 3M | 6M | 12M |
| --- | --- | --- | --- | --- | --- | --- | --- | --- | --- | --- |
| 11 | 67149640 | KDM2A | 113 | 22 | 143856 | 3715 | KDM2A; ADRBK1; ANKRD13D | FALSE | FALSE | TRUE |
| 11 | 66130442 | PACS1 | 92 | 13 | 172960 | 2405 | PACS1; GAL3ST3; SF3B2 | FALSE | TRUE | TRUE |
| 17 | 78020335 | TNRC6C | 85 | 29 | 143507 | 1807 | TNRC6C; TMC6; TMC8; C17orf99 | FALSE | FALSE | FALSE |
| 17 | 31250620 | NF1 | 88 | 4 | 293588 | 1289 | NF1; MIR4733 | FALSE | FALSE | TRUE |
| 11 | 65458065 | MALAT1 | 69 | 6 | 118661 | 1142 | MALAT1; FRMD8; NEAT1; MIR612; SCYL1; LTBP3 | FALSE | FALSE | FALSE |
| 17 | 81583163 | NPLOC4 | 53 | 1 | 125601 | 1015 | NPLOC4; ACTG1; C17orf70; TSPAN10; OXLD1; CCDC137 | FALSE | TRUE | TRUE |
| 11 | 73071469 | FCHSD2 | 77 | 1 | 343225 | 821 | FCHSD2; STARD10 | FALSE | FALSE | TRUE |
| 22 | 50354381 | PPP6R2 | 48 | 1 | 151125 | 777 | PPP6R2; PLXNB2; DENND6B; SBF1 | FALSE | FALSE | FALSE |
| 3 | 47083604 | SETD2 | 52 | 6 | 167118 | 770 | SETD2; NBEAL2; KIF9-AS1 | FALSE | FALSE | TRUE |
| 16 | 68127705 | NFATC3 | 48 | 1 | 178999 | 710 | NFATC3; DUS2; SLC7A6 | FALSE | FALSE | FALSE |

#### 3.2.3 Risk Gene Screening

Risk gene screening (100 kb distance threshold, 0.1% clone contribution threshold) identified 23 integration sites near AE-associated genes, including 4 sites within 10 kb of LMO2, 3 sites near MECOM, and 2 sites near CCND2. Cancer gene screening revealed 187 integration sites near oncogenes or tumor suppressors, with 12 high-abundance clones (>1% contribution) requiring enhanced monitoring. Immune gene screening identified 412 integration sites near immune-related genes, predominantly in patients with higher vector copy numbers. These findings demonstrate LISA’s utility in systematic safety assessment and risk stratification.

#### 3.2.4 Longitudinal Clonal Dynamics

Longitudinal analysis revealed stable polyclonal hematopoiesis across the 18-month follow-up period. Richness increased slightly from M6 to M18 (10.8 → 11.3, log_2_ scale), indicating continued detection of low-abundance clones. Evenness remained relatively stable (8.2 → 8.5), and PMD showed no significant trend (−2.6 → −2.8), suggesting maintenance of polyclonal diversity without emergence of dominant clones. The dominant clone early warning system flagged 3 integration sites with sustained increasing abundance trends (Mann-Kendall p < 0.05), all below the 1% threshold and not near risk genes, indicating acceptable safety profiles. These results demonstrate LISA’s capability for comprehensive longitudinal monitoring and early detection of potential safety signals.

### 3.3 Comparison with Existing Tools

We compared LISA’s functionality with existing integration site analysis tools (Table 4). ISAnalytics provides basic integration site processing but lacks Safe Harbor annotation, risk gene screening, and dominant clone warning. MELISSA focuses on clonal reconstruction from co-occurrence patterns rather than comprehensive integration site characterization. MultIS specializes in clonal tracking but does not detect CIS or perform functional annotation. LISA is the only tool providing: (1) optimized CIS detection with scalable algorithm, (2) systematic Safe Harbor and risk gene annotation, (3) comprehensive longitudinal analysis with PMD calculation, (4) dominant clone early warning system, and (5) complete set of publication-ready visualizations. The unique combination of efficiency, comprehensiveness, and clinical relevance positions LISA as the most suitable tool for gene therapy safety assessment.

**Table 4.** Functional comparison of LISA with existing integration site analysis tools

| Feature | LISA | ISAnalytics | MELISSA | MultIS |
| --- | --- | --- | --- | --- |
| **CIS Detection** | ✓ ($O(n \log n)$) | ✓ | ✓ | ✗ |
| **Safe Harbor Annotation** | ✓ | ✗ | ✗ | ✗ |
| **Enhancer/Promoter Annotation** | ✓ | Partial | ✗ | ✗ |
| **Risk Gene Screening** | ✓ (3 gene sets) | ✗ | ✗ | ✗ |
| **AE Gene Screening** | ✓ (36 genes) | ✗ | ✗ | ✗ |
| **Cancer Gene Screening** | ✓ (980 genes) | ✗ | ✗ | ✗ |
| **Longitudinal PMD Analysis** | ✓ | Partial | ✗ | ✗ |
| **Dominant Clone Warning** | ✓ | ✗ | ✗ | ✗ |
| **Interactive Visualization** | ✓ (6 types) | Partial | ✓ | ✓ |
| **Algorithm Complexity** | $O(n \log n)$ | Not specified | Not specified | $O(n^2)$ |

## 4. Discussion

We developed LISA as a comprehensive solution for integration site analysis specifically tailored to gene therapy safety assessment. Three key innovations distinguish LISA from existing tools: algorithmic optimization enabling analysis of large-scale clinical datasets, systematic safety-focused functional annotation including Safe Harbor regions and curated risk gene sets, and comprehensive longitudinal monitoring with early warning for dominant clones.

The algorithmic innovations (chromosome-position encoding and seed-site strategy) reduce CIS detection complexity from O(n^2^) to O(n log n), enabling real-time analysis of large datasets. This performance improvement is critical for clinical applications where rapid turnaround supports timely decision-making. The scalability to >100,000 integration sites positions LISA for large-scale post-market surveillance and meta-analyses across multiple trials.

LISA’s functional annotation system directly addresses regulatory requirements for gene therapy safety evaluation. Safe Harbor identification helps assess integration patterns relative to low-risk genomic regions. The three curated risk gene sets (AE, cancer, immune) enable systematic screening for integrations near genes implicated in adverse events or malignant transformation. This risk-stratified approach supports both regulatory submissions and clinical decision-making by highlighting integration sites warranting enhanced monitoring.

For longitudinal studies, LISA’s PMD calculation and dominant clone early warning system provide quantitative metrics for tracking clonal dynamics. Decreasing PMD or emergence of dominant clones may signal loss of polyclonality preceding clinical events. Early detection of such signals could enable proactive monitoring or intervention before safety concerns become clinically manifest. The statistical framework (Mann-Kendall trend test) provides objective, reproducible assessment of clonal trends.

Several limitations should be noted. First, LISA currently supports only hg38 human reference genome; extension to other species would require additional reference data curation. Second, Safe Harbor definitions are based on current literature and may require periodic updates as knowledge evolves. Third, the dominant clone warning model requires validation with larger datasets including confirmed adverse events to calibrate sensitivity and specificity. Fourth, LISA analyzes integration site distributions but does not assess clone functionality (e.g., differentiation capacity, proliferation rate); integration with transcriptomic or epigenomic data would enhance risk assessment.

Future development directions include: (1) integration with single-cell sequencing data for lineage tracing, (2) machine learning models predicting integration site risk based on multi-omic features, (3) expansion to include mutation detection at integration sites, (4) development of a web server interface for broader accessibility, and (5) establishment of a community database for integration site data sharing and meta-analysis. These enhancements will further strengthen LISA’s utility in advancing gene therapy safety science.

In conclusion, LISA addresses a critical gap in the gene therapy safety assessment toolkit by providing an efficient, comprehensive, and clinically relevant platform for integration site analysis. The package’s alignment with regulatory requirements, validation with real clinical data, and unique combination of features position it as a valuable resource for gene therapy development, clinical monitoring, and post-market surveillance. By enabling systematic, quantitative safety assessment, LISA contributes to the continued advancement of gene therapy as a safe and effective therapeutic modality.

## 5. Availability and Implementation

LISA is implemented as an R package requiring R version 4.0 or higher. Key dependencies include GenomicRanges, TxDb.Hsapiens.UCSC.hg38.knownGene, org.Hs.eg.db, ggplot2, ggrepel, ggpubr, patchwork, visNetwork, igraph, RIdeogram, and readxl. The package is freely available at [GitHub repository URL to be added] under the MIT open-source license. Comprehensive documentation includes function references, a detailed vignette demonstrating the complete analysis workflow, and example datasets in the inst/extdata directory. Installation instructions, tutorials, and updates are available at the project website.

## Acknowledgments Funding

This work was supported by grants from the National Natural Science Foundation of China (No.82330005 to HM, No.82070124 to HM), and the National Science Fund for Distinguished Young Scholars (No. 82425003 to HM).

